# Photoplethysmography-based glucose estimation with feature selection

**DOI:** 10.64898/2026.10.07.757242

**Authors:** Swandip Singha, Aditta Chowdhury

**Affiliations:** Department of Electrical and Electronic Engineering, Chittagong University of Engineering and Technology, Chattogram, Bangladesh

**Keywords:** Photoplethysmography (PPG), Non-invasive glucose estimation, Feature selection, Ensemble machine learning, Signal quality index, Diabetes screening

## Abstract

Diabetes mellitus remains a major global health challenge, while conventional blood glucose monitoring relies on invasive fingerstick methods that limit continuous and large-scale screening. This study investigates non-invasive glucose estimation using photoplethysmography (PPG) signals processed through an adaptive digital filtering pipeline combined with a multi-criteria statistical feature selection framework. A custom acquisition system was used to collect PPG recordings from 51 subjects, from which 14 morphological and autonomic features spanning time-domain, derivative, frequency-domain, and nonlinear characteristics were extracted. Feature relevance was evaluated using Pearson and Spearman correlations, partial correlation with confounder adjustment, mutual information, and composite ranking. The validated pipeline was subsequently applied to a public four-wavelength PPG benchmark dataset comprising 133 subjects, evaluating three progressively enriched machine learning input configurations. The proposed approach, combining subject metadata with greedy forward-selected PPG features, achieved an *R*^2^ of 0.410 using Random Forest at the 730 nm channel, outperforming the metadata-only baseline by 0.055. All configurations maintained 100% Clarke Error Grid Zone A+B compliance, confirming clinical safety. These findings demonstrate that a compact and interpretable set of PPG-derived autonomic and morphological features can significantly enhance demographic predictors for non-invasive glucose estimation.

## 1 Introduction

Diabetes mellitus remains one of the most consequential chronic diseases of the modern era, placing enormous strain on healthcare systems across both high-income and resource-limited settings. The International Diabetes Federation estimates that approximately 537 million adults were affected globally in 2021, with projections indicating this number will exceed 783 million by 2045 absent meaningful intervention [1]. Poorly controlled glycaemia drives cardiovascular complications, nephropathy, retinopathy, and peripheral neuropathy, all of which are substantially preventable through early detection and consistent monitoring [2]. Despite this burden, blood glucose assessment still relies predominantly on capillary sampling via enzymatic test strips — an inherently invasive, episodic method that discourages the frequent measurement effective diabetes management demands and renders population-scale screening impractical in resource-constrained settings.

Photoplethysmography (PPG) is among the most promising non-invasive candidates for continuous glycaemic monitoring. By detecting volumetric changes in peripheral blood flow through optical absorption at one or more wavelengths, a PPG sensor captures haemodynamic information about arterial compliance, cardiac timing, and autonomic vascular regulation [3]. Chronic hyperglycaemia compromises endothelial function, accelerates arterial stiffening, and induces autonomic neuropathy — physiological changes that manifest as measurable alterations in PPG morphology, spectral content, and nonlinear dynamics [4, 5]. Multi-wavelength PPG systems extend this capability further by simultaneously probing tissue at several spectral bands, enabling differential haemoglobin absorption to be exploited for richer haemodynamic characterisation from a single compact front-end [6]. The ubiquity of PPG sensors in consumer smartwatches and pulse oximeters positions this modality as uniquely suited to low-cost, point-of-care deployment.

Machine learning has become central to wearable physiological sensing, enabling clinically relevant quantities to be extracted from complex, noisy biosignals. Ensemble methods — particularly Random Forest and Extremely Randomized Trees — are well-suited to moderate-dimensional biomedical feature sets owing to their tolerance of feature collinearity, resistance to overfitting, and natural capacity for predictor importance estimation [7, 8]. Support vector regression with nonlinear kernels provides a competitive alternative on small datasets where margin maximisation offers generalisability benefits [9]. Deep convolutional and recurrent architectures have also been applied to raw PPG waveforms for blood pressure and glucose prediction; however, their data demands and limited physiological interpretability constrain their utility where sample sizes are small and explainability is required [10]. Across all paradigms, principled feature engineering grounded in signal physiology is as determinative of final accuracy as model architecture [11].

Several structural limitations persist in published work on PPG-based glucose estimation: feature-glucose correlations are reported without adjusting for haemodynamic confounders [12, 13]; signal quality assessment is rarely incorporated upstream of feature extraction [14]; studies are typically confined to a single platform/wavelength, preventing cross-dataset generalisability assessment [15, 16]; feature selection is often reduced to univariate filter ranking without cross-validated confirmation [17]; and the incremental value of PPG features over demographic/haematological metadata is seldom quantified [18].

This paper presents a two-stage framework for PPG-based glucose estimation. First, a custom PPG acquisition system collected beat-level recordings from 51 subjects using a three-criterion artifact rejection scheme and a composite signal quality index to retain only high-fidelity segments. Fourteen physiologically motivated features (time-domain, derivative VPG/APG, frequency-domain, and non-linear), plus four engineered composite indices, were extracted and ranked with a multi-criteria statistical framework (Pearson/Spearman correlation, partial correlation with confounder control, mutual information, and a weighted composite score) to identify glucose-relevant, confounder-robust predictors. Second, the top-ranked features were evaluated on a four-wavelength public PPG benchmark under three progressively enriched machine learning input configurations, showing consistent improvement over baseline models while maintaining 100% Clarke Error Grid Zone A+B compliance and ISO 15197 standards.

## 2 Literature Review

### A. Non-Invasive Modalities and Physiological Basis

Optical non-invasive glucose monitoring spans several spectral modalities. NIR spectroscopy exploits glucose absorption in the 700–2500 nm window but suffers calibration drift from inter-subject skin and temperature variability [15]; Mid-infrared spectroscopy offers stronger specificity but is impractical for wearables [16]; Raman spectroscopy and optical coherence tomography show bench-level sensitivity but require hardware incompatible with mass-market devices [19]; and indirect methods (pulse transit time, skin impedance, sweat metabolites) remain limited by intra-subject variability and poor specificity [17]. PPG stands out for its commodity hardware, low power, and ability to encode multiple cardiovascular and autonomic parameters from a single optical front-end [3, 4]. Its waveform morphology encodes arterial compliance and peripheral resistance through the systolic upstroke, dicrotic notch, and diastolic decay — parameters progressively impaired in diabetic vasculopathy [5]. The acceleration photoplethysmogram (APG), the second derivative of the PPG, amplifies these distinctions; its early systolic peak-to-trough ratio is a validated surrogate of vascular age [20]. High-frequency spectral power (0.15–0.4 Hz) reflects parasympathetic cardiac modulation linked to insulin resistance [4], while Poincaré SD1/SD2 decompose RR-interval variability into components sensitive to autonomic neuropathy [21]. Multi-wavelength acquisition further expands the feature space via differential haemoglobin absorption across spectral channels [6].

### B. Machine Learning and Signal Quality Assessment

Supervised ensemble regression dominates PPG-based glucose prediction. Random Forest [7] and Extremely Randomized Trees [8] handle feature collinearity, resist overfitting, and provide built-in importance rankings; compact, curated input sets consistently outperform full unselected feature vectors [18]. SVR with RBF kernels generalises reliably on small biomedical cohorts [9, 12], gradient-boosted methods offer additional regularisation but require careful tuning on limited datasets [22], and deep learning on raw PPG sequences shows promise for blood pressure estimation, yet data demands and opacity to clinical interpretation restrict its glucose-prediction applicability [10, 23]. Upstream of modelling, motion artifacts, baseline wander, and irregular inter-beat intervals can silently corrupt extracted features unless explicitly gated [14]. Zero-phase Butterworth and FIR bandpass filters suppress out-of-band noise while preserving morphologically critical waveform landmarks [3], and composite signal quality indices (SQIs) — aggregating inter-beat timing consistency, amplitude regularity, and template correlation — provide principled, data-driven exclusion of low quality segments [14, 24]. Despite evidence that artifact contaminated beats degrade prediction accuracy by margins comparable to model-selection effects, formal SQI gating remains largely absent from published PPG-based glucose estimation workflows [14].

### C. Feature Selection and Confounder Control

Effective feature selection for glucose estimation must distinguish genuine glucose associations from confounder-mediated correlations with blood pressure and heart rate [12, 13]. Partial correlation, which residualises both feature and target against the confounder matrix before measuring association, provides a tractable correction [11]. Mutual information captures nonlinear dependencies complementary to Pearson and Spearman metrics; integrating all three into a composite ranking score enables physiologically grounded shortlisting [25], and greedy forward selection with cross-validated evaluation then yields a compact subset free from the optimistic bias of filter-only methods [18]. Although each technique has been validated individually, their integration into a unified confounderaware pipeline for PPG-based glucose prediction — with formal SQI gating and explicit quantification of the marginal value of PPG features over demographic/haematological metadata — has not previously been reported. The present framework addresses this gap through composite SQI gating, multicriteria confounder-adjusted feature ranking, cross-validated forward selection, and three-configuration evaluation on a four-wavelength public benchmark.

## 3 Data Acquisition and Dataset Description

Two datasets were employed in this study: a locally collected dataset, used for feature extraction and selection to identify the most informative features, and a publicly available benchmark 4-wavelength PPG dataset [26], subsequently used to evaluate the generalizability of the selected features and the proposed approach.

### 3.0.1 Primary Dataset: Custom PPG Acquisition System

The primary dataset was gathered using a low-cost, custom-built acquisition system comprising an Arduino Uno microcontroller interfaced with a MAX30100 pulse oximeter sensor, as illustrated in Fig. 1. The MAX30100 operates in reflective mode, acquiring the PPG signal at the fingertip through integrated infrared and red LED. Reference measurements of SBP and DBP were recorded simultaneously using a calibrated digital sphygmomanometer, while blood glucose concentration was obtained via a standard glucometer immediately prior to signal acquisition. All measurements were stored in CSV format for subsequent processing.

**Fig. 1.**
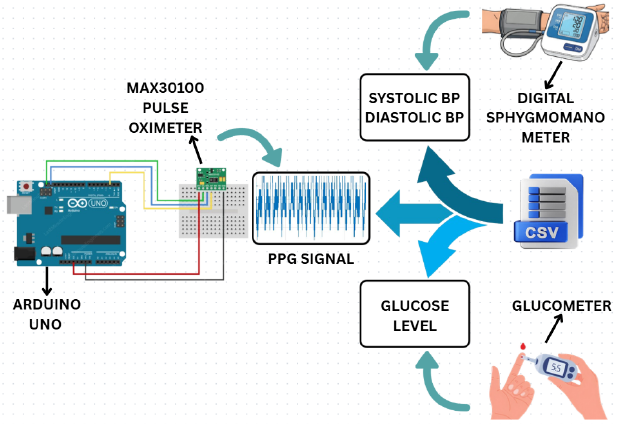
PPG data acquisition setup. The MAX30100 pulse oximeter, interfaced with an Arduino Uno, captures the raw PPG signal.

The primary dataset encompasses recordings from 51 subjects and four target physiological variables: systolic blood pressure (SBP), diastolic blood pressure (DBP), heart rate (HR), and blood glucose. The signals were sampled at 125 Hz (8 ms interval), with each recording lasting approximately 4.8 s. Descriptive statistics are presented in Table 1.

**Table 1.** Descriptive statistics of the four target physiological variables in the primary dataset.

| Variable | Unit | Min | Max | Variance |
| --- | --- | --- | --- | --- |
| Systolic BP | mmHg | 79.0 | 138.0 | 161.73 |
| Diastolic BP | mmHg | 45.0 | 88.0 | 72.70 |
| Heart Rate | bpm | 54.0 | 115.0 | 127.69 |
| Blood Glucose | mmol/L | 4.8 | 6.8 | 0.22 |

### 3.0.2 Secondary Dataset: Four-Wavelength PPG Benchmark

To assess cross-dataset generalisability, experiments were also conducted on a publicly available benchmark dataset: the Blood Pressure Measurement based on Four-Wavelength PPG Signals dataset. Signals were acquired at the index finger in reflective mode across four wavelength channels: 660 nm, 730 nm, 850 nm, and 940 nm. Reference SBP and DBP values were measured immediately before signal collection using an OMRON HEM7201 oscillometric device. The dataset contains recordings from 180 subjects, each comprising 60 seconds of continuous PPG signal sampled at 200 Hz, along with subject-level metadata including SBP and DBP.

For cross-dataset evaluation, the PPG morphological features extracted from the primary dataset pipeline were re-derived from the secondary dataset’s signals, enabling a direct comparison of model performance across acquisition platforms and population groups.

## 4 Methodology

The proposed methodology is organised into two self-contained but conceptually aligned workflows. The first is applied to the custom-built primary dataset: it encompasses signal preprocessing, beat-level PPG feature extraction, and a statistical feature-analysis stage that identifies physiologically meaningful, glucose-relevant features. The second applies the validated processing pipeline to the four-wavelength public benchmark dataset and evaluates three machine learning input configurations for blood glucose prediction.

### 4.1 Primary Dataset: Signal Processing, Feature Extraction, and Statistical Analysis

The primary pipeline, illustrated in Fig. 2, receives the raw PPG signal from the MAX30100 acquisition system and passes it through four sequential stages, described below.

**Fig. 2.**
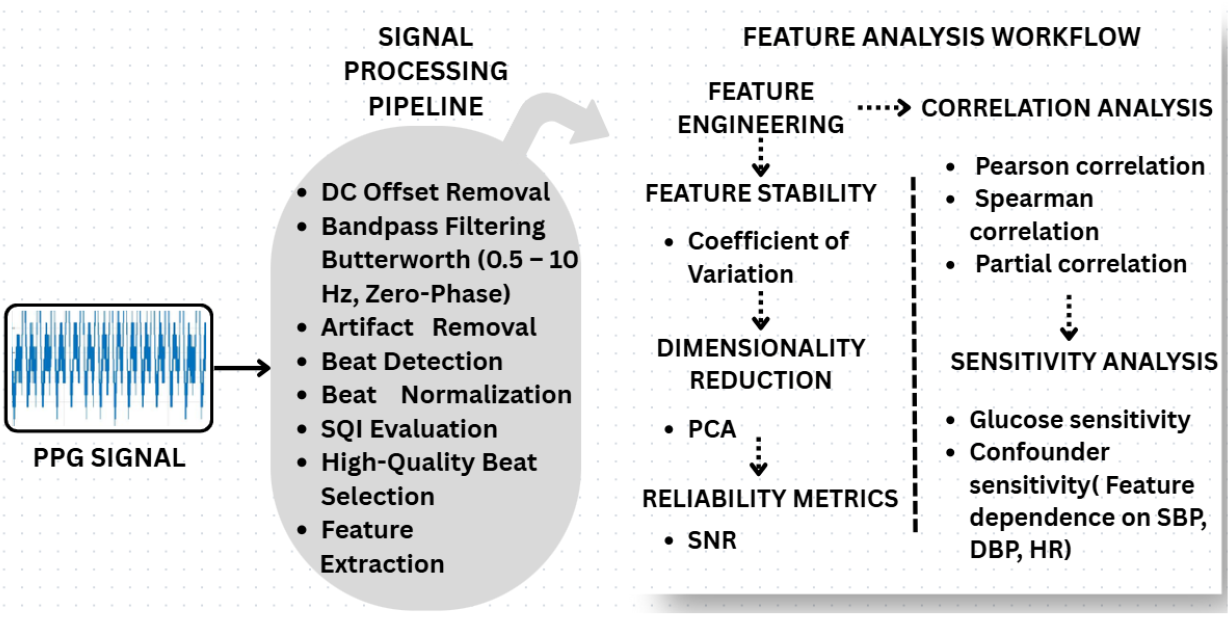
Overview of the primary dataset methodology, including signal processing and feature analysis workflows.

#### 4.1.1 Signal Preprocessing

Each raw PPG recording sampled at *f*_*s*_ = 125 Hz is first DC-corrected by subtracting its mean:

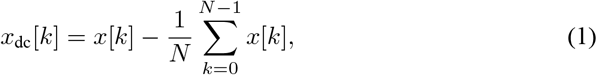

where *x*[*k*] is the raw signal and *N* is the number of samples. The DC-corrected signal is bandpass filtered using a zero-phase, 4thorder Butterworth filter with passband [0.5, 10] Hz to suppress baseline wander and high-frequency noise while preserving the dicrotic notch and morphological harmonics:

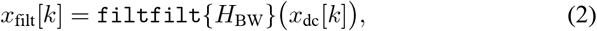

where *H*_BW_ has the squared magnitude response

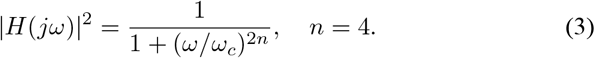

Zerophase implementation prevents group-delay distortion, preserving the timing of morphology-critical features.

Artifact-contaminated samples are identified from *x*_filt_[*k*] using three criteria:

1. Amplitude criterion: a sample is flagged if

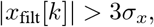

where *σ*_*x*_ is the standard deviation of the filtered signal.
2. Gradient criterion: a sample is flagged if

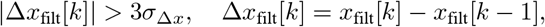

where *σ*_Δ*x*_ is the standard deviation of the gradient.
3. RR interval criterion: a beat is flagged if either

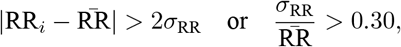

where RR_*i*_ is the *i*-th inter-beat interval, 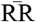 is the mean RR interval, and *σ*_RR_ is the standard deviation. Samples between consecutive beats are flagged if the first condition is met, and the entire signal is flagged if the coefficient of variation exceeds 0.30.

Flagged samples, along with a symmetric 10-sample buffer, are replaced with `NaN` and excluded from all downstream calculations.

#### 4.1.2 Beat Detection, Normalization, and SQI Evaluation

Systolic peaks are identified on the cleaned signal using a prominence-based detector with a minimum inter-peak distance of 0.4 s (corresponding to a physiological ceiling of 150 bpm) and a normalised amplitude threshold of 0.30. Only peaks whose inter-beat intervals fall within [0.4, 1.5] s are retained. Each consecutive beat segment *B*_*i*_ = *x*[*k*_*i*_ : *k*_*i*+1_] is resampled to a fixed length of 100 points via cubic-spline interpolation:

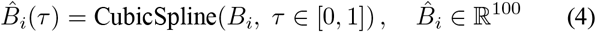

enabling cross-subject morphological comparison regardless of instantaneous heart rate.

A composite SQI is computed for each beat from three sub-indices. The RR consistency index penalises irregular inter-beat timing:

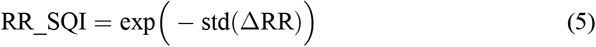

The amplitude consistency index penalises beat-to-beat amplitude variation:

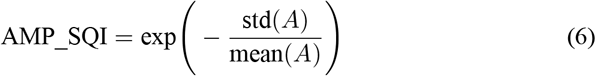

The morphological consistency index quantifies waveform regularity via correlation with the median template beat 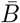:

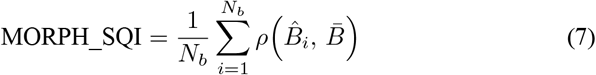

where *ρ*(*·*) denotes the Pearson correlation coefficient and *N*_*b*_ is the number of beats. The final composite SQI is the unweighted mean:

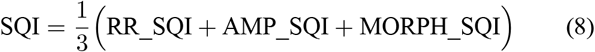

A data-driven exclusion threshold, set at the 20th percentile of all observed beat-level SQI scores, is applied; beats below this threshold are discarded. The SQI distribution across the 51 subjects ranged from 0.780 to 0.995 (Fig. 3), confirming high overall recording quality. After applying artifact and SQI criteria, 32 subjects were retained for further analysis.

**Fig. 3.**
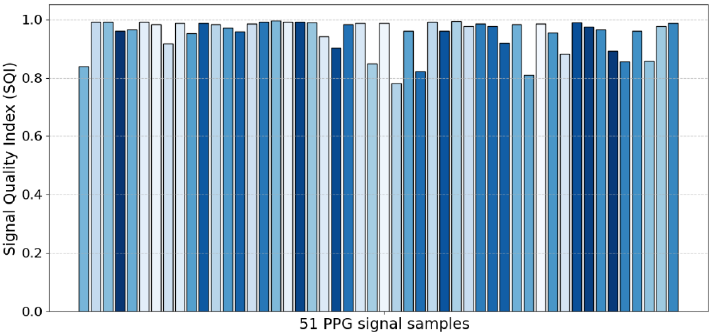
SQI of the best 600sample segment for each of the 51 PPG signal samples in the primary dataset.

#### 4.1.3 Beat-Level Feature Extraction

Following quality-controlled beat selection, 14 features are extracted per subject across four physiologically motivated categories (Table 2; landmarks annotated in Fig. 4).

**Table 2.**
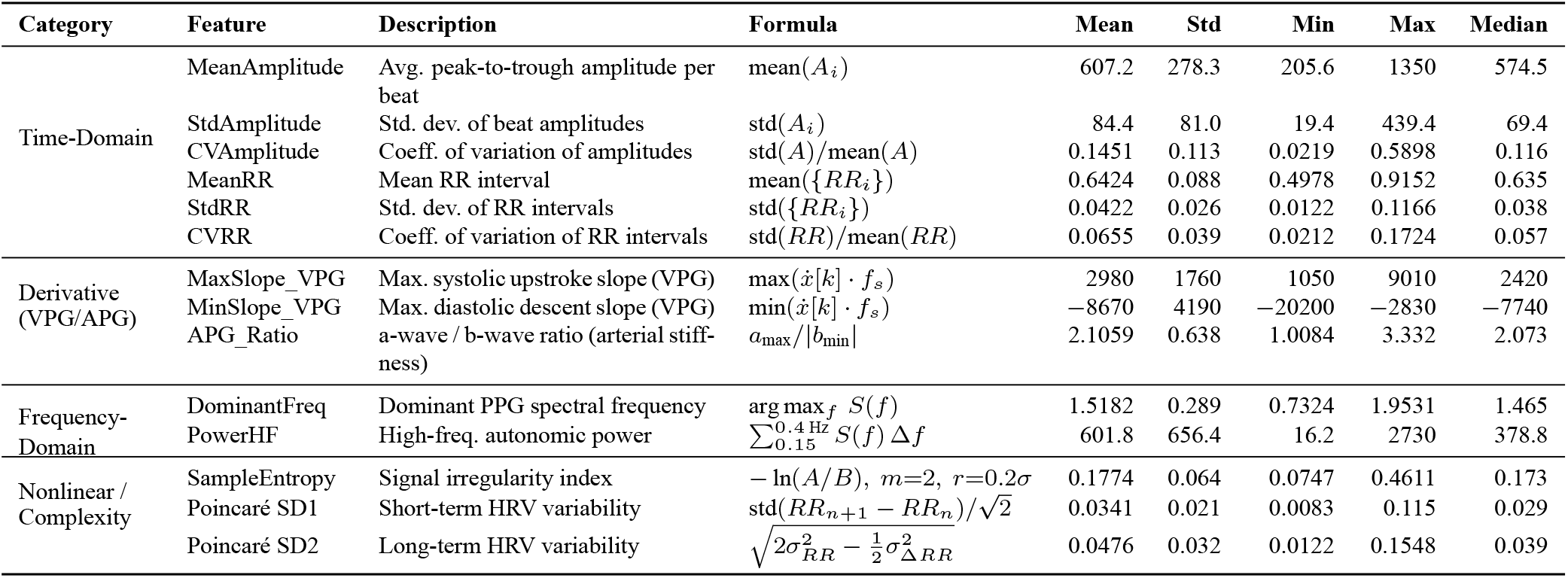
Taxonomy of the 14 extracted PPG features computed from the primary dataset.

**Fig. 4.**
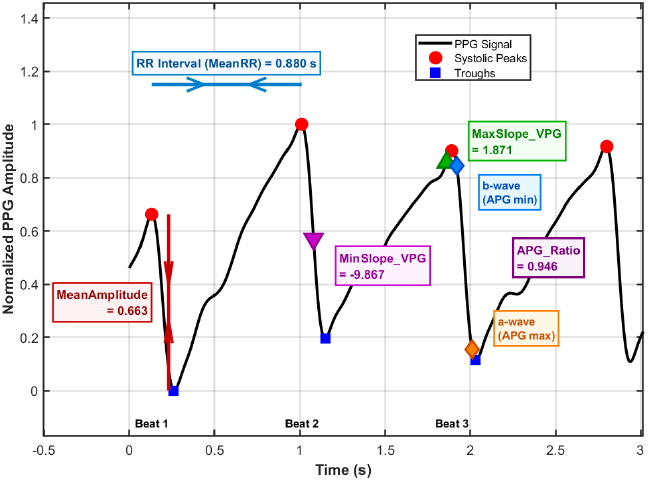
Annotation of extracted morphological features on a representative normalised PPG waveform.

1. Time-domain features: For each beat *i*, the amplitude 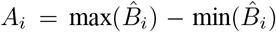 is computed. MeanAmpli-tude and StdAmplitude characterise average pulse magnitude and beat-to-beat variability, while CVAmplitude normalises variability by the mean. The RR-interval features (MeanRR, StdRR, CVRR) quantify cardiac timing regularity, reflecting autonomic modulation of vascular tone — a pathway linking the autonomic nervous system to glucose homeostasis.
2. Derivative-based features (VPG and APG): The velocity photoplethysmogram (VPG) is the first temporal derivative of the filtered signal, 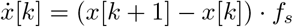. MaxSlope_VPG captures the peak rate of systolic pressure rise, MinSlope_VPG the diastolic runoff rate. The APG (second derivative, 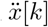) yields APG_Ratio, the ratio of the first positive peak (a-wave) to the first negative peak (bwave), a well-established surrogate for arterial wall stiffness:

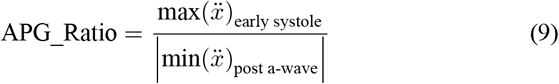
3. Frequency-domain features: The power spectral density *S*(*f*) is estimated via Welch’s method (256-point Hamming window, 50% overlap). Dominant-Freq identifies the cardiac fundamental frequency; PowerHF (0.15–0.4 Hz) captures parasympathetic autonomic spectral contribution.
4. Nonlinear features: Sample entropy quantifies signal irregularity (*m* = 2, *r* = 0.2 *σ*_*x*_):

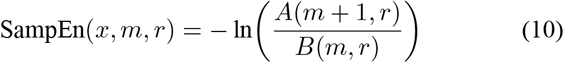

where *B*(*m, r*) and *A*(*m* + 1, *r*) count matched template pairs of length *m* and *m* + 1. Poincaré SD1/SD2 decompose RR-interval variability into short- and long-term components:

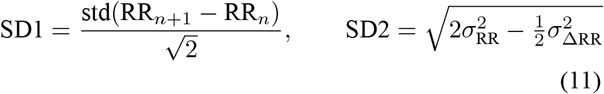

#### 4.1.4 Feature Engineering and Statistical Analysis

Four composite features were derived to capture non-additive physiological interactions not represented in the base set:

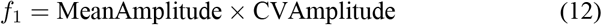

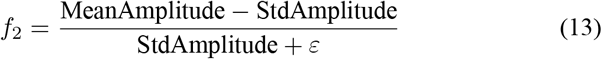

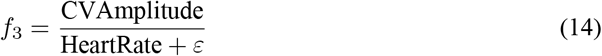

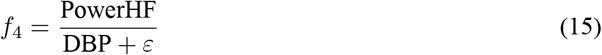

where *ε* = 0.001 prevents division by zero.

Statistical analyses were conducted to assess feature relevance and robustness with respect to blood glucose:

1. Feature relevance: Pearson (*r*) and Spearman (*ρ*_*s*_) correlations with *p*-values quantify linear and monotonic associations between each feature *X* and glucose *Y*.
2. Partial correlation with confounder control: controlling for SBP, DBP, and HR, both *X* and *Y* were regressed on the confounder matrix **Z**, and correlation was computed between residuals:

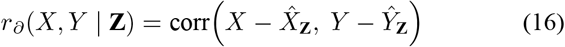
3. Sensitivity analysis: glucose sensitivity via standardised regression coefficients (*β*) on *z*-scored features; Confounder sensitivity via *R*^2^ between each feature and SBP, DBP, HR.
4. Non-linear dependency: mutual information regression captures nonlinear feature–glucose dependencies.
5. Dimensionality reduction: PCA on the *z*-scored feature matrix estimates intrinsic dimensionality.
6. Composite feature ranking: relevance, robustness, and stability were integrated into

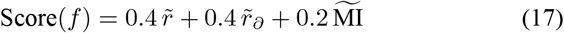

with all quantities min–max normalised to [0, 1]; top-ranked features were selected for further evaluation.

### 4.2 Secondary Dataset: Feature Re-derivation and Machine Learning Evaluation

The secondary pipeline (Fig. 5) applies the validated processing and feature extraction stages to the fourwavelength public PPG benchmark and evaluates blood glucose predicttion across three ML input configurations.

**Fig. 5.**
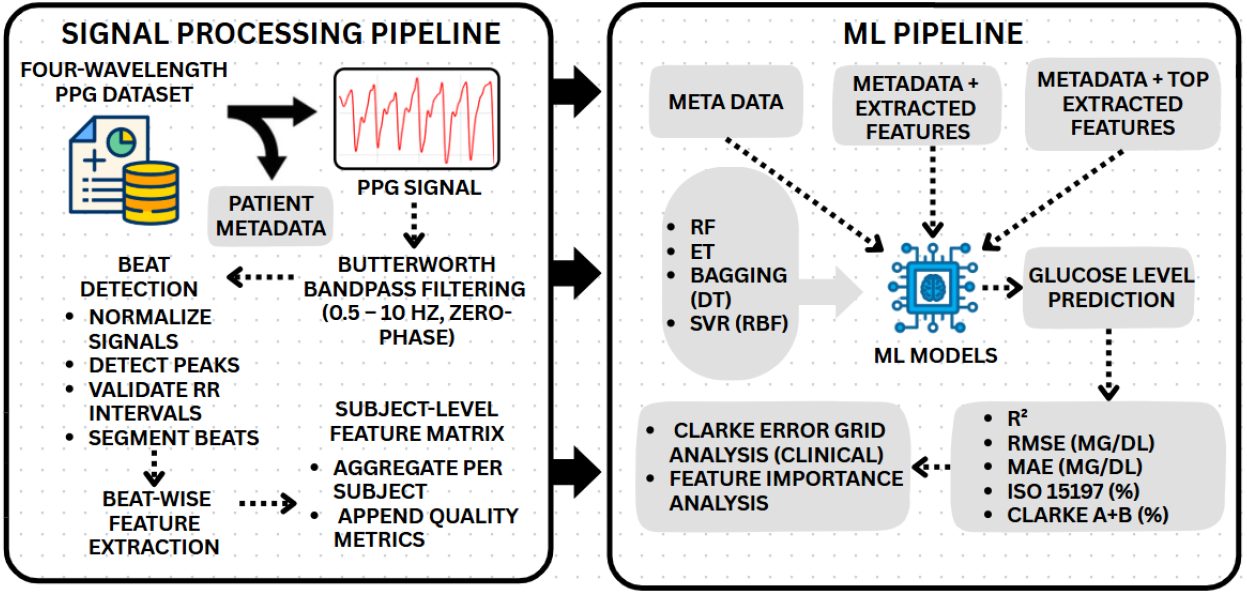
Secondary dataset signal processing and machine-learning pipeline.

#### 4.2.1 Signal Processing for the Four-Wavelength Dataset

The four-wavelength dataset (660, 730, 850, 940 nm) is sampled at *f*_*s*_ = 200 Hz. An identical signal processing pipeline is applied independently to each wavelength channel. A linear-phase FIR bandpass filter (order 200, Hamming window, passband [0.4, 12] Hz) improves harmonic preservation at the higher sampling rate while ensuring zero-phase distortion. Peak detection follows the prominence-based scheme of Section 4.1.2, with minimum inter-peak distance rescaled to 0.4 *× f*_*s*_ = 80 samples. Beat-wise feature extraction follows the 14-feature taxonomy of Table 2, with the four engineered features of Section 4.1.4 computed per recording.

From the original cohort of 180 subjects, 133 with complete physiological metadata (gender, age, height, weight, haemoglobin, SBP, DBP) were retained for model training and evaluation; metadata were incorporated at the modelling stage rather than during signal preprocessing.

#### 4.2.2 Machine Learning Input Configurations

Three progressively enriched input configurations were evaluated:

1. *Configuration 1*: metadata only (gender, age, height, weight, haemoglobin, SBP, DBP) — the baseline.
2. *Configuration 2*: metadata plus all extracted PPG features (14 primary + 4 engineered), a 25-dimensional input vector.
3. *Configuration 3*: metadata plus a subset of top PPG features selected via greedy forward selection, iteratively choosing features that yield the largest cross-validated *R*^2^ improvement (typically top 3–4 features).

All input features are scaled using a `RobustScaler`:

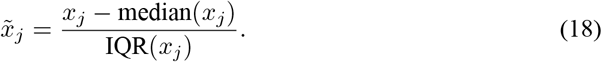

#### 4.2.3 Regression Models and Hyperparameter Optimisation

Four regression models were trained per configuration: Random Forest (RF) and Extra Trees (ET), bagging-based ensembles built on bootstrapped subsets and feature-random subspaces; Bagging with Decision Tree (DT) base estimators as a baseline ensemble comparator; and Support Vector Regression with RBF kernel (SVR-RBF):

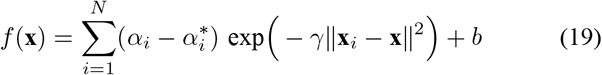

Hyperparameter optimisation used 5-fold cross-validated grid search on the training set; the held-out test set comprised 20% of subjects, stratified by glucose quartile bins.

#### 4.2.4 Evaluation Metrics

1. Standard metrics: *R*^2^, RMSE, and MAE, computed in mg/dL (mmol/L *×* 18.0182).
2. ISO 15197:2013 compliance: a prediction *ĝ* is compliant if

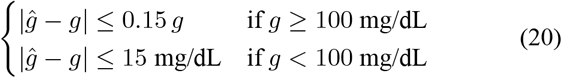

where *g* is the reference glucometer value.
3. Clarke Error Grid Analysis (CEGA): classifies each prediction–reference pair into clinical zones; Zone A is clinically accurate (*±*20%), Zone B is benign, and Zones C–E denote increasingly dangerous misclassification. The combined Zone A+B rate is the primary safety metric:

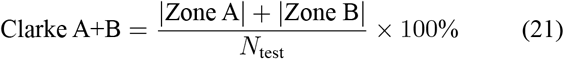
4. Feature importance: mean impurity decrease (Gini importance) for tree-based models, applied to the best-performing model per configuration.

## 5 Results

This section reports the primary dataset signal-processing and feature-analysis outcomes, the secondary dataset preprocessing/filter performance, and blood glucose prediction across the three machine learning configurations.

### 5.1 Primary Dataset Results

#### 5.1.1 PPG Signal Quality and Beat Detection

Table 3 summarises signal quality and beat detection statistics. Of the 51 subjects recruited, 32 met the quality criteria after applying the composite SQI threshold and artifact rejection scheme, yielding 95 valid beats for feature extraction. Across the full cohort, 364 beats were initially detected, with 2.97 *±* 2.16 valid beats per subject. Filtered-signal heart rate averaged 95.06 *±* 12.53 bpm (mean RR 0.642*±*0.088 s), and motion artifact contamination remained modest (9.64 *±* 4.01% mean rate).

**Table 3.** PPG signal-level statistics for the primary dataset.

| Category | Metric | Mean ± Std | Range |
| --- | --- | --- | --- |
| Beat Segmentation | Total beats detected | 364 | – |
| | Mean valid beats per subject | $2.97 \pm 2.16$ | 1 – 9 |
|  | Median valid beats per subject | 2 | – |
| Heart Rate | HR (bpm) | $95.06 \pm 12.53$ | 65.56 – 120.54 |
| | RR interval (s) | $0.642 \pm 0.088$ | – |
| Artifact Rate | Motion artifact (%) | $9.64 \pm 4.01$ | 0 – 15.33 |

The processing pipeline for a representative subject is illustrated in Fig. 6: the raw signal (top-left) shows a periodic pulsatile pattern with baseline drift; after zero-phase Butterworth filtering the systolic peak, dicrotic notch, and diastolic phase become clearly identifiable; four systolic peaks are annotated after artifact removal and peak detection, with a mean beat-level SQI of 0.904 against a data-driven threshold of 0.722. The right panel shows normalised beat morphology for the same subject, with individual beats aligned and overlaid on the median template, demonstrating high morphological consistency.

**Fig. 6.**
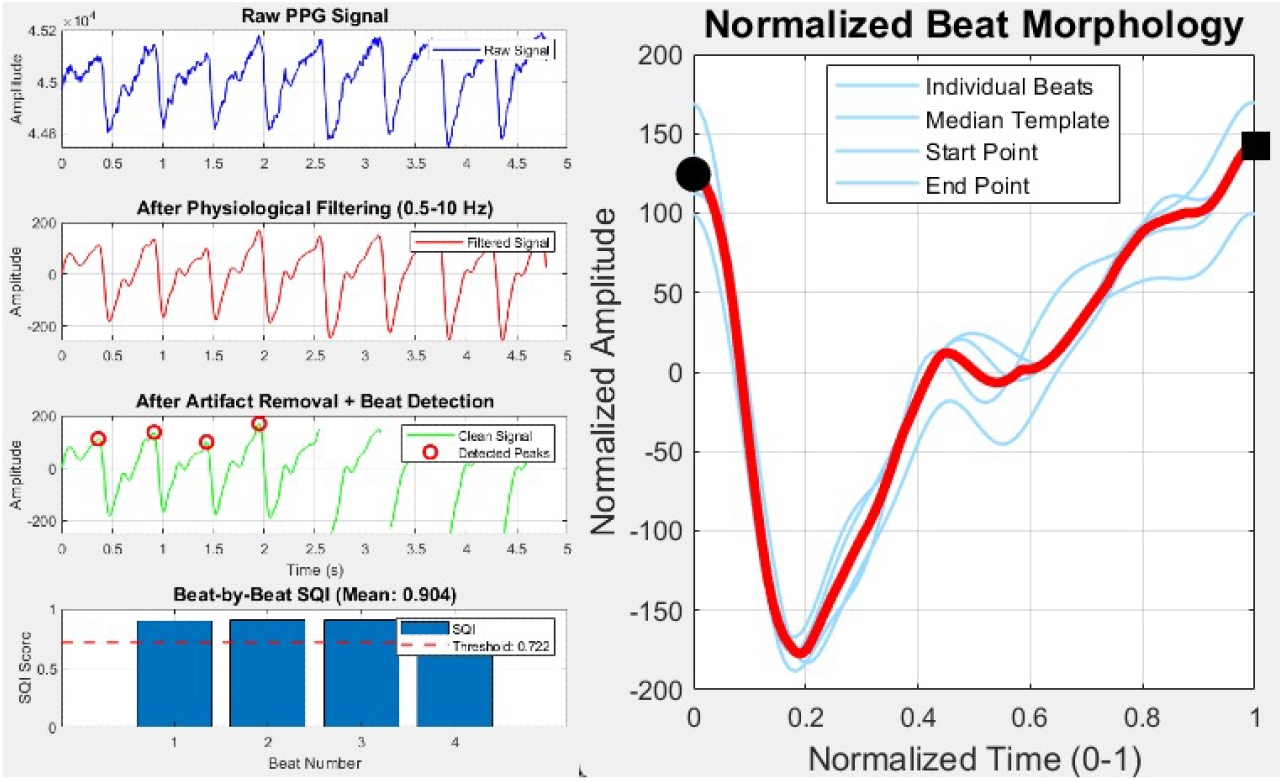
PPG signal processing pipeline with filtering, SQI assessment, and beat morphology analysis.

#### 5.1.2 Feature Analysis and Selection

Statistical feature analysis was performed on the complete feature set; after composite score–based ranking, the top eight features were selected (Fig. 7).

**Fig. 7.**
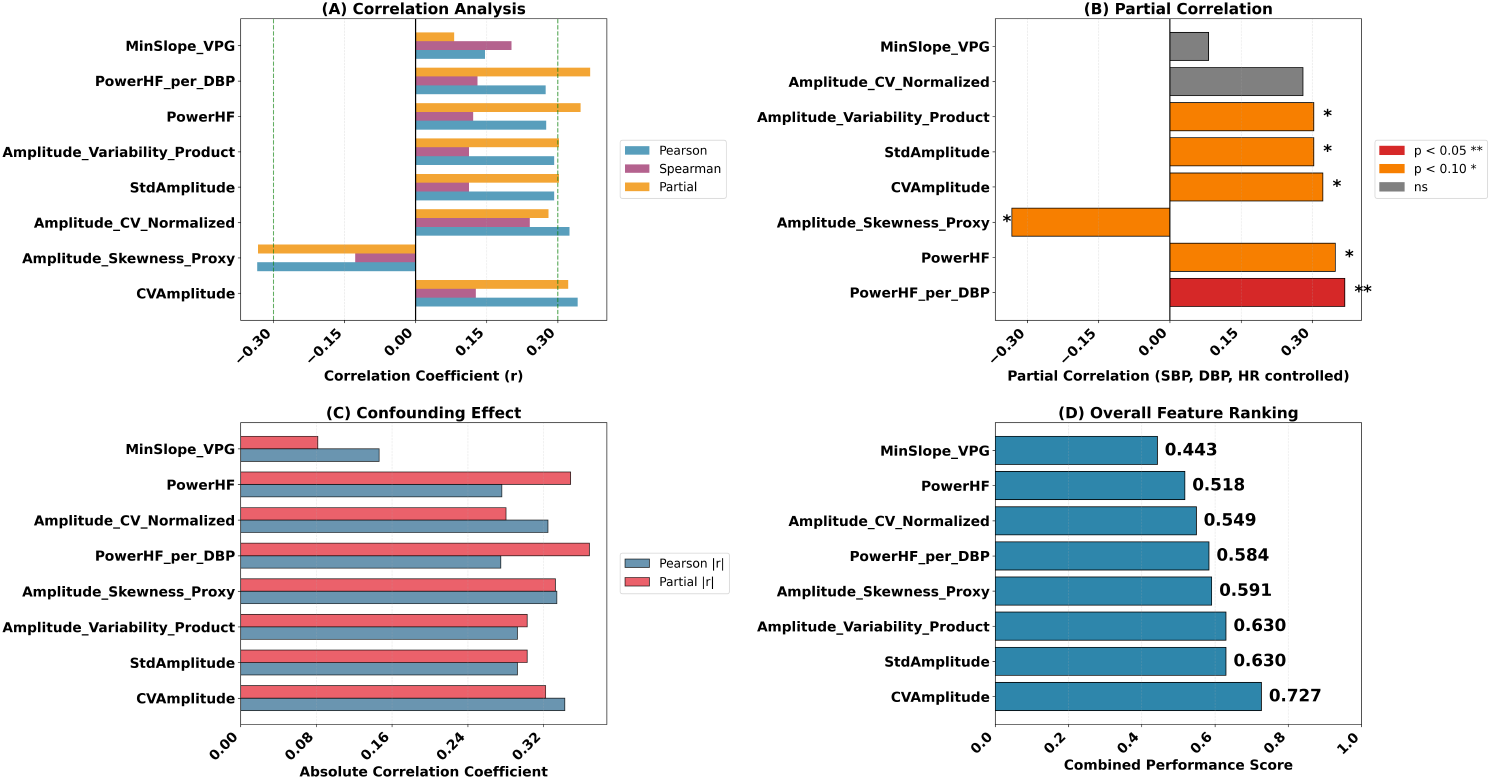
Statistical feature analysis: (A) Pearson, Spearman, and partial correlations with blood glucose; (B) partial correlations after confounder control; (C) confounding effect comparison; (D) composite feature ranking score.

1. Correlation analysis: CVAmplitude exhibited the highest positive correlation across Pearson, Spearman, and partial measures (Pearson *r ≈* 0.30; partial *r*_*∂*_ *≈* 0.30), robust to SBP/DBP/HR adjustment. Amplitude_Skewness_Proxy showed a negative partial correlation (*≈ −*0.30), and PowerHF_per_DBP showed the largest partial correlation among engineered features (*r*_*∂*_ *≈* 0.35). Pearson and Spearman correlations agreed directionally for the top features, indicating predominantly monotonic relationships.
2. Significance: PowerHF_per_DBP was significant at *p <* 0.05; CVAmplitude and PowerHF reached *p <* 0.10; Amplitude_Skewness_Proxy was significant at *p <* 0.05 (negative); MinSlope_VPG was non-significant.
3. Sensitivity and confounder analysis: Table 4 reports standardised glucose sensitivity (*β*) and confounder *R*^2^ values. CVAmplitude showed the largest *β* = 0.3424 with negligible confounder dependence 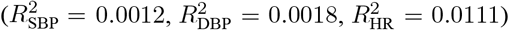, confirming it as the most glucoses-pecific feature. Amplitude_Skewness_Proxy showed *β* = *−* 0.3342 with moderate HR dependence 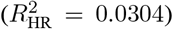, and PowerHF_per_DBP showed 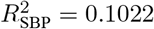 (expected, as DBP appears in its denominator). Most Pearson correlations were preserved after partial adjustment, with only modest reductions.
4. Mutual information: MI values were generally low, indicating weak non-linear feature–glucose dependence, consistent with the correlation-based analysis; MI served as a complementary rather than primary selection criterion.
5. Dimensionality analysis: PCA showed high feature redundancy — the first two components captured 59.4% of variance, and seven components explained 95%, supporting the use of aggregated/composite representations.
6. Composite ranking: CVAmplitude achieved the highest composite score (0.727), confirming it as the most informative glucose-relevant feature.

**Table 4.** Sensitivity analysis for the top-ranked PPG features: standardised glucose sensitivity coefficient (*β*) and confounder *R*^2^ values with respect to SBP, DBP, and HR.

| Feature | Pearson $p$ | Spearman $p$ | $\beta$ (Glucose) | $R^2$ (SBP) | $R^2$ (DBP) | $R^2$ (HR) |
| --- | --- | --- | --- | --- | --- | --- |
| CVAmplitude | 0.0551 | 0.4875 | +0.3424 | 0.0012 | 0.0018 | 0.0111 |
| StdAmplitude | 0.1044 | 0.5389 | +0.2924 | 0.0115 | 0.0003 | 0.0017 |
| Amplitude_Variability_Product | 0.1044 | 0.5389 | +0.2924 | 0.0115 | 0.0003 | 0.0017 |
| Amplitude_Skewness_Proxy | 0.0616 | 0.4875 | −0.3342 | 0.0038 | 0.0129 | 0.0304 |
| PowerHF_per_DBP | 0.1279 | 0.4755 | +0.2748 | 0.1022 | 0.0307 | 0.0014 |
| Amplitude_CV_Normalized | 0.0698 | 0.1840 | +0.3247 | 0.0004 | 0.0000 | 0.0694 |
| PowerHF | 0.1263 | 0.5081 | +0.2759 | 0.0605 | 0.0060 | 0.0038 |
| MinSlope_VPG | 0.4239 | 0.2665 | +0.1464 | 0.0155 | 0.0001 | 0.0776 |

### 5.2 Secondary Dataset Results

#### 5.2.1 Four-Wavelength PPG Preprocessing

The linear-phase FIR bandpass filter was applied independently to each wavelength channel. Fig. 8 shows raw and filtered spectra: the filter suppresses content below 0.4 Hz and above 12 Hz while preserving cardiac spectral content, with the characteristic FIR stop-band ripple visible beyond 12 Hz, confirming correct operation.

**Fig. 8.**
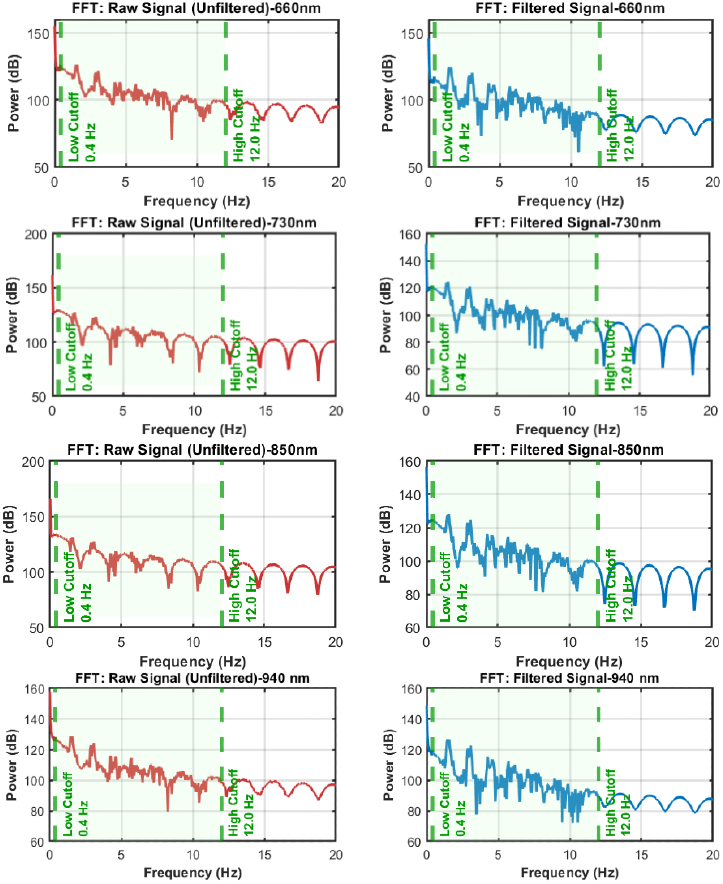
Frequency spectra of raw (red) and filtered (blue) PPG signals for all four wavelength channels (660, 730, 850, and 940 nm).

Table 5 summarises filter performance: SNR improved across all channels; baseline wander power was reduced by 10^15^–10^16^ and high-frequency noise (*>*10 Hz) by *∼* 10^12^ in every channel. Signal preservation (1–4 Hz band) ranged from 43.17% (730 nm) to 69.27% (850 nm), reflecting aggressive attenuation of out-of-band harmonics, while morphology correlation (0.022–0.109) reflects the elimination of low-frequency baseline components that dominated the raw signals.

**Table 5.** FIR bandpass filter performance metrics for each of the four wavelength channels in the secondary dataset. Changes are reported relative to the unfiltered raw signal.

| Metric | 660 nm | 730 nm | 850 nm | 940 nm |
| --- | --- | --- | --- | --- |
| SNR (dB) | +2.82 (increase) | +5.40 (increase) | +2.52 (increase) | +5.38 (increase) |
| Baseline Power (<0.3 Hz) | $1.29 \times 10^{16}$ (decrease) | $2.91 \times 10^{15}$ (decrease) | $3.49 \times 10^{16}$ (decrease) | $5.86 \times 10^{15}$ (decrease) |
| High Freq. Noise Power (>10 Hz) | $3.65 \times 10^{12}$ (decrease) | $1.05 \times 10^{12}$ (decrease) | $9.65 \times 10^{12}$ (decrease) | $1.92 \times 10^{12}$ (decrease) |
| PPG Signal Power (1–4 Hz) | $3.60 \times 10^{13}$ (decrease) | $1.21 \times 10^{13}$ (decrease) | $9.39 \times 10^{13}$ (decrease) | $2.14 \times 10^{13}$ (decrease) |
| Signal Preservation (%) | 66.73 | 43.17 | 69.27 | 56.42 |
| Morphology Correlation | 0.022 | 0.102 | 0.0399 | 0.109 |

#### 5.2.2 Machine Learning Results

Table 6 reports the performance of all evaluated models across the three configurations (for Configurations 2 and 3, only the best-performing model per wavelength is shown for clarity).

**Table 6.** Comparison of model performance across three configurations on the secondary dataset.

| Configuration | Wavelength | Model | $R^2$ | RMSE (mg/dL) | MAE (mg/dL) | ISO 15197 (%) |
| --- | --- | --- | --- | --- | --- | --- |
| Config-1 |  | **Random Forest** | **0.355** | **12.69** | **9.56** | **74.07** |
|  |  | Extra Trees | 0.313 | 13.09 | 8.91 | 81.48 |
|  |  | Bagging (Trees) | 0.321 | 13.01 | 9.56 | 74.07 |
|  |  | SVR (RBF) | 0.316 | 13.06 | 8.96 | 81.48 |
| Config-2 | 660 nm | Random Forest | 0.244 | 13.73 | 10.10 | 81.48 |
|  | 730 nm | Random Forest | 0.278 | 13.42 | 10.04 | 77.78 |
|  | 850 nm | Extra Trees | 0.252 | 13.65 | 10.03 | 81.48 |
|  | 940 nm | Bagging (Trees) | 0.153 | 14.53 | 11.30 | 77.78 |
| Config-3 | 660 nm | Random Forest | 0.379 | 12.44 | 8.77 | 77.78 |
|  | 730 nm | Random Forest | 0.410 | 12.13 | 8.98 | 81.48 |
|  | 850 nm | Random Forest | 0.378 | 12.45 | 9.65 | 81.48 |
|  | 940 nm | SVR (RBF) | 0.353 | 12.70 | 9.03 | 85.19 |

##### Configuration 1: Metadata Only (Baseline)

Random Forest achieved the highest *R*^2^ = 0.355; Extra Trees, Bagging, and SVR-RBF attained comparable *R*^2^ in the range 0.313–0.321.

Fig. 9 shows the feature importance ranking of the best-performing Random Forest model alongside its Clarke Error Grid. Haemoglobin (g/L) emerged as the most important predictor (*≈* 0.25), followed by age, SBP, height, and DBP; gender contributed negligibly (*<*0.01). All test-set predictions fell within Zone A or Zone B, with none in the dangerous Zones C–E; predictions clustered near 80–100 mg/dL, reflecting the narrow glucose distribution of the cohort (4.8–6.8 mmol/L; 86.5–122.5 mg/dL). All four models achieved 100% Zone A+B compliance, with ISO 15197 compliance ranging from 74.07% to 81.48%.

**Fig. 9.**
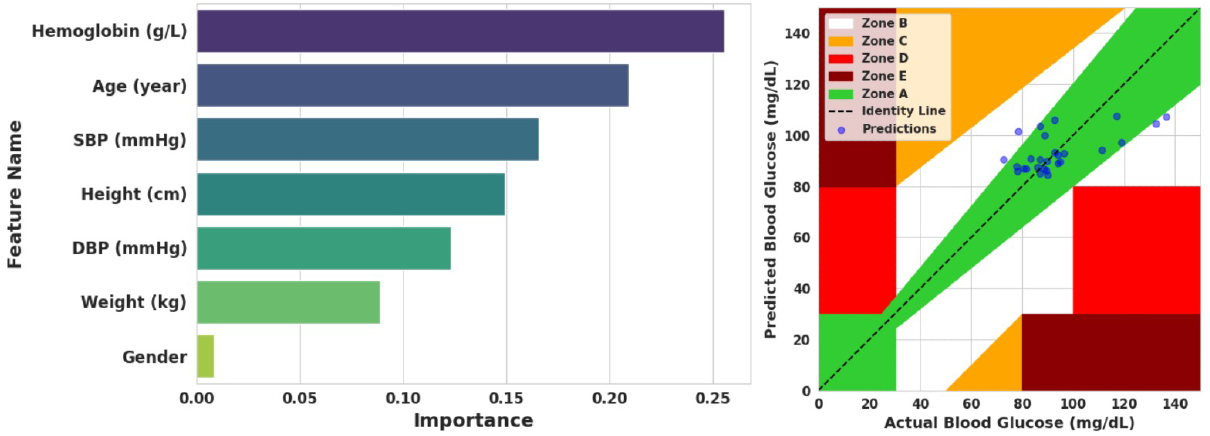
Configuration 1 results for the best model (RF).

##### Configuration 2: Metadata Plus All Extracted PPG Features

Compared with Configuration 1 (*R*^2^ = 0.355), Configuration 2 showed no improvement at any wavelength; the full 18feature set likely introduced redundancy and variance without proportionate bias reduction, and wavelength dependent SNR differences further reduced feature effectiveness. The 730 nm channel performed best among wavelengths, though still below baseline. Fig. 10 shows the top-10 importances per channel: haemoglobin and age consistently ranked highest, and several PPG-derived features appeared in the top-10 without improving overall performance.

**Fig. 10.**
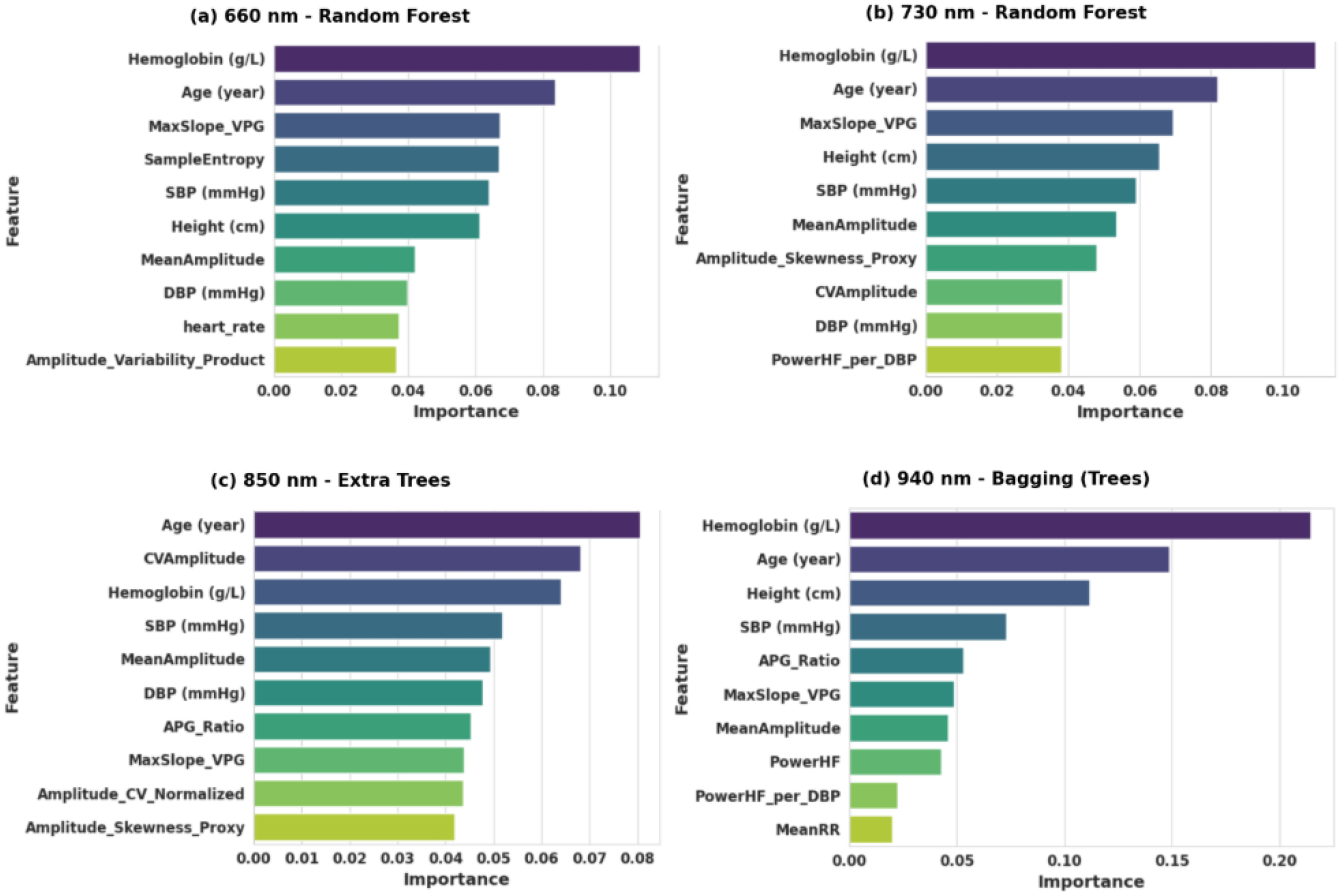
Top-10 feature importances for Configuration 2 across four wavelength channels.

##### Configuration 3: Metadata Plus Selected Top PPG Features

Table 7 lists the greedy forwardselected features per wave-length; this configuration produced the strongest overall results, with all channels achieving *R*^2^ *>* 0.35 and 100% Clarke A+B compliance. The 730 nm Random Forest model achieved the best overall performance (*R*^2^ = 0.410), followed by 660 nm and 850 nm (*R*^2^ = 0.379 and 0.378); the 940 nm SVR-RBF model achieved *R*^2^ = 0.353 and the highest ISO 15197 compliance (85.19%). Relative to Configuration 2, selected features yielded consistent gains of 0.08–0.13 in *R*^2^ and reductions of 0.8–1.8 mg/dL in RMSE, confirming that feature selection improves generalisability by removing noise from irrelevant/redundant inputs.

**Table 7.** Wavelength-wise features selected via greedy forward feature selection for Configuration 3.

| Wavelength | Selected Features |
| --- | --- |
| 660 nm | PowerHF per DBP, Amplitude Variability Product, PowerHF, Amplitude Skewness Proxy |
| 730 nm | StdAmplitude, PowerHF per DBP, Amplitude Variability Product, Amplitude CV Normalized |
| 850 nm | CVAmplitude, PowerHF, Amplitude Skewness Proxy, PowerHF per DBP, StdAmplitude, Amplitude Variability Product, Amplitude CV Normalized |
| 940 nm | PowerHF, Amplitude CV Normalized, Amplitude Variability Product, StdAmplitude, PowerHF per DBP |

Fig. 11 shows top-10 importances for the tree-based channels (660, 730, 850 nm; SVR-RBF at 940 nm has no tree-based importances). Age and haemoglobin consistently dominated; among PPG-derived features, amplitude variability metrics (Amplitude Variability Product, CVAmplitude) were the most informative, with PowerHF and Pow erHF_per_DBP also contributing — supporting the physiological link between autonomic PPG features and glucose via sympathetic–parasympathetic balance and peripheral vascular resistance.

**Fig. 11.**
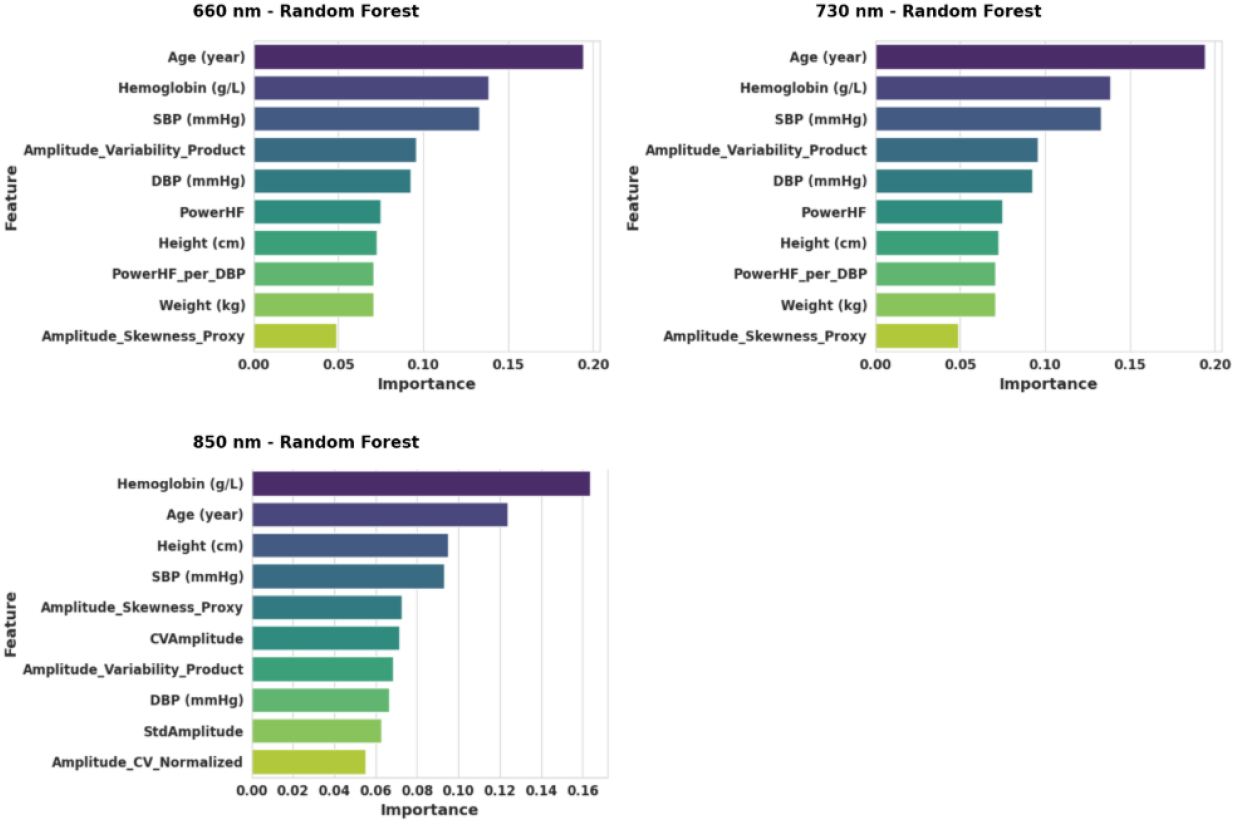
Top-10 feature importances for Configuration 3 for the three wavelength channels with treebased best models.

Fig. 12 shows the Clarke Error Grid for the best model per wavelength: all four channels maintained 100% Clarke A+B compliance with no predictions in Zones C–E. The 660 nm and 730 nm Random Forest models produced the tightest clusters along the identity line, consistent with their higher *R*^2^; the 850 nm scatter is slightly broader and shifted toward 80–95 mg/dL, while the 940 nm SVRRBF model spans 80–110 mg/dL with a modest upward bias. Predictions across all channels remain concentrated in the 80–105 mg/dL range, reflecting the narrow glucose distribution of the 133 retained subjects.

**Fig. 12.**
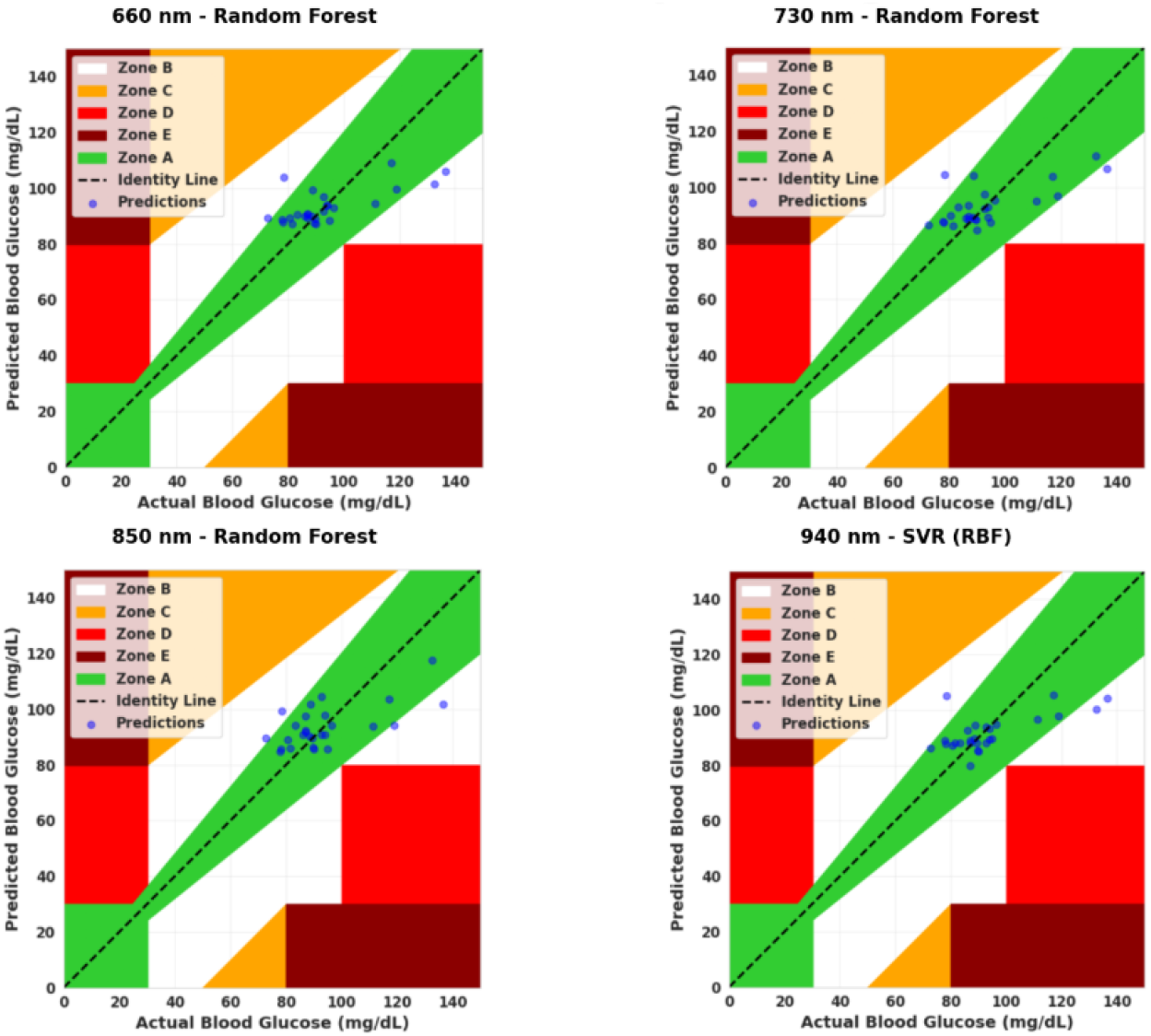
Clarke Error Grid Analysis for Configuration 3.

Configuration 3 consistently outperformed both the metadataonly baseline and full-feature augmentation across all four wavelengths, establishing that a compact set of carefully selected PPG morphological and autonomic features gives the best tradeoff between accuracy and complexity. The 730 nm channel was optimal, with Random Forest attaining *R*^2^ = 0.410 — the highest recorded in the study, a gain of 0.055 over Configuration 1 (*R*^2^ = 0.355) achieved through only four added PPG features. The degradation in Configuration 2 relative to Configuration 1 is attributed to redundancy and noise from including all 18 PPG features without selection.

## 6 Conclusion

This study introduced a structured two-stage framework for non-invasive blood glucose estimation using photoplethysmography. A zero-phase adaptive bandpass filtering pipeline combined with composite signal quality assessment enabled reliable extraction of high-quality beat segments. From these signals, 14 physiologically meaningful features were derived and ranked through a multi-criteria statistical evaluation strategy. Experiments on a four-wavelength public benchmark showed that selectively augmenting subject metadata with three to four top-ranked PPG features consistently improved predictive performance over both metadata-only and full-feature configurations. The best performance was obtained at 730 nm with an R^2^ of 0.410 while maintaining 100% Clarke Error Grid Zone A+B compliance across all wavelength channels. Amplitude variability metrics and parasympathetic spectral indicators (HF-band power) provided the most consistent physiological signals associated with glucose variation after confounder adjustment. These findings support the feasibility of low-cost wearable PPG-based systems for large-scale diabetes screening. Future work should focus on larger datasets, modeling intra-day glucose variability, and deep learning approaches for end-to-end physiological representation learning.

## Author contributions

Swandip Singha: Methodology, conceptualization, visualization, writing original draft

Aditta Chowdhury: Supervision, methodology, writing original draft

## Funding

The authors have no relevant financial or non-financial interests to disclose.

## Competing Interests

The authors have no competing interests to declare that are relevant to the content of this article.

## Data Availability Statement

Datasets used in this study are mentioned and cited within the manuscript.

## Consent to Publish declaration

not applicable

## Consent to Participate declaration

not applicable

## Ethics declaration

not applicable

## Notes

### Competing Interest Statement

The authors have declared no competing interest.

### Summary of Updates

The manuscript has been revised to improve organization, clarity, figure presentation, and consistency throughout the text and supplementary materials.

